# Functional connectome contractions in temporal lobe epilepsy: microstructural underpinnings and associations to surgical outcome

**DOI:** 10.1101/756494

**Authors:** Sara Larivière, Yifei Weng, Reinder Vos de Wael, Birgit Frauscher, Zhengge Wang, Andrea Bernasconi, Neda Bernasconi, Dewi V. Schrader, Zhiqiang Zhang, Boris C. Bernhardt

**Author notes:** authors contributed equally (Zhiqiang Zhang, Boris C. Bernhardt). **Correspondence to:** Boris C. Bernhardt, PhD, Montreal Neurological Institute (NW-256), 3801 University Street, Montreal, Quebec, Canada H3A 2B4, Sara Larivière, PhD Candidate.

## Abstract

Temporal lobe epilepsy (TLE) is the most common drug-resistant epilepsy in adults. While commonly related to hippocampal pathology, increasing evidence suggests structural changes beyond the mesiotemporal lobe. Functional anomalies and their link to underlying structural alterations, however, remain incompletely understood. We studied 30 drug-resistant TLE patients and 57 healthy controls using multimodal magnetic resonance imaging analyses. We developed a novel framework that parameterizes functional connectivity distance, consolidating functional and geometric properties of macroscale networks. Compared to controls, TLE showed connectivity distance reductions in temporo-insular and prefrontal networks, suggesting topological segregation of functional networks. Our novel approach furthermore allowed for the testing of morphological and microstructural associations, and revealed that functional connectivity contractions occurred independently from TLE-related cortical atrophy but were mediated by microstructural changes in the underlying white matter. All patients underwent a comparable resective surgery after our study and a regularized supervised machine learning paradigm with 5-fold cross-validation demonstrated that patient-specific functional anomalies predicted post-surgical seizure outcome with 74±8% accuracy, outperforming classifiers operating on clinical and structural imaging features. Our findings suggest connectivity distance contractions as a clinically relevant pathoconnectomic substrate of TLE. Functional topological isolation may represent a microstructurally mediated network mechanism that tilts the balance towards epileptogenesis.

## Introduction

Temporal lobe epilepsy (TLE) is a recognized surgically-amenable disorder and ranks among the most prevalent drug-resistant adult epilepsies (Wiebe et al. 2001). While traditionally conceptualized as a prototypical “focal” epilepsy with a mesiotemporal epicenter, mounting histopathological and neuroimaging evidence has shown structural and functional compromise affecting widely distributed brain networks, supporting the notion of TLE as a system disorder affecting large-scale networks (Gleichgerrcht et al. 2015; Tavakol et al. 2019). In effect, adequately capturing its connectopathy may advance our understanding of brain mechanisms giving rise to temporal lobe seizures, improve diagnostic procedures of individual patients, and guide surgical prognostics.

Paralleling prior studies mapping whole-brain structural alterations in TLE via volumetric (Lin et al. 2007; McDonald et al. 2008; Bernhardt et al. 2010) and diffusion MRI techniques (Bonilha et al. 2013; Liu et al. 2016), recent years have witnessed a surge in resting-state functional MRI (rs-fMRI) analyses (Liao et al. 2010; He et al. 2015). Given its versatility to probe multiple functional systems within a single acquisition, rs-fMRI represents a candidate technique to identify network mechanisms and biomarkers of brain disorders. Initial rs-fMRI investigations have targeted connectivity alterations in TLE compared to controls in specific networks, showing mainly decreased connectivity when focussing on mesiotemporal circuits (Morgan et al. 2011), on the interplay of mesial and lateral temporal regions (Maccotta et al. 2013), or when studying connectivity between temporal seeds and macroscale networks, particularly the so-called “default-mode” network (Bernhardt, Bernasconi, Liu, et al. 2016).

The focus on individual regions or networks has precluded an integrative view of functional network anomalies in TLE, and particularly their associations to underlying perturbations of cortical morphology and microstructure. To fill this gap, we developed a novel rs-fMRI paradigm that systematically profiled functional connections across the entire cortical mantle. Our approach combined conventional functional connectomics with a brain-wide geodesic distance mapping technique to estimate connectivity length distributions of any given cortical area, and thus, draw specific inferences on the balance of short- and long-range connectivity. Specifically, we tested the hypothesis that aberrant local connectivity and reduced long-range connections may form to topologically isolated functional networks that contribute to recurrent seizure activity. Such mechanism would be in line with findings from experimental models of limbic epilepsy, which posit that aberrant local connectivity and deafferentation contribute to connectivity contractions, which in turn may ultimately promote recurrent seizure activity (Knopp et al. 2005; Sharma et al. 2007). To assess structural underpinnings of these functional connectome reconfigurations, we further examined associations between connectivity distance shifts and MRI measures of cortical morphology and white matter microstructure. Finally, as all patients underwent anterior temporal lobectomy after our imaging investigations, we harnessed machine learning techniques and cross-referenced pre-surgical connectome features with individualized resection cavity data to identify regional predictors or post-surgical outcome prognosis.

## Materials and Methods

### Participants and multimodal MRI dataset

We studied a consecutive cohort of 30 drug-resistant unilateral TLE patients (15 males, mean±SD=26.9±8.7 years, 19 right-sided focus). Patients were diagnosed according to the classification of the International League Against Epilepsy (ILAE) based on a comprehensive examination that includes clinical history, seizure semiology, continuous video-EEG telemetry recordings, neuroimaging, and neuropsychology. Patients had a mean±SD duration of epilepsy of 11.4±7.9 years (range=2–356 months). No patient had encephalitis, malformations of cortical development, (*e.g.*, tumors, vascular malformations), or a history of traumatic brain injury.

Patients were selected from a larger patient cohort and met the following inclusion criteria: (*i*) patients underwent anterior temporal lobectomy as a treatment of their seizures at Jinling Hospital between August 2008 to April 2017 (mean±SD=4.3±5.7 months, range=0.1–29 months after our imaging investigations), (*ii*) patients had a research-dedicated high-resolution 3T MRI before surgery that included anatomical (*T1w*), functional (*rs-fMRI*), and diffusion (*dMRI*) imaging, (*iii*) patients had post-operative histological confirmation of hippocampal sclerosis, (*iv*) patients had post-surgical imaging to confirm the location of the surgical cavity, and (*v*) post-surgical seizure outcome was available and determined according to Engel’s modified classification (Engel 1993) with a follow-up of at least one year after surgery (mean±SD=3.23±2.4 years, range=1-9 years). Twenty patients (66%) were seizure-free (Engel-I), while four (17%) showed significant reductions in seizure frequency (Engel-II), four (17%) showed prolonged seizure-free intervals (Engel-III), and two showed no worthwhile improvement (Engel-IV). Detailed information on the patient cohort is presented in **Supplementary table 1**.

Patients were compared to 57 age- and sex-matched healthy individuals (25 males, mean±SD=25.6±5.9 years) who underwent identical imaging.

This study was approved by Research Ethics Board of Jinling Hospital, Nanjing University School of Medicine, and written informed consent was obtained from all participants.

### MRI acquisition

Multimodal MRI data were obtained on a Siemens Trio 3T scanner (Siemens, Erlangen, Germany) and included: (*i*) high-resolution 3D T1-weighted MRI using a magnetization-prepared rapid gradient-echo sequence (*T1w*, repetition time [TR] = 2300 ms, echo time [TE] = 2.98 ms, flip angle = 9º, voxel size = 0.5 × 0.5 × 1 mm^3^, field of view [FOV] = 256 × 256 mm^2^, 176 slices), (*ii*) resting-state blood-oxygen-level dependent (BOLD) functional MRI using a single-shot, gradient-recalled echo planar imaging sequence (*rs-fMRI*, 255 volumes, TR = 2000 ms, TE = 30 ms, flip angle = 90º, FOV = 240 × 240 mm^2^, voxel size = 3.75 × 3.75 × 4 mm^3^, 30 slices), and (*iii*) diffusion MRI using a spin echo-based echo planar imaging sequence with 4 b0 images (*dwi*, TR = 6100 ms, TE = 93 ms, flip angle = 90º, FOV = 240 × 240 mm^2^, voxel size = 0.94 × 0.94 × 3 mm^3^, b-value = 1000 s/mm^2^, diffusion directions = 120). During the rs-fMRI acquisition, participants were instructed to keep their eyes closed, to not think of anything in particular, and to stay awake. While every participant underwent T1w and rs-fMRI scans, only a subset of controls underwent diffusion MRI (31/57 controls, 14 males, mean±SD age: 27.3±7.4 years; 30/30 TLE patients). To increase power, the main functional analyses were carried out by comparing the TLE patients against all (*i.e.*, 57/57) controls, whereas analyses involving diffusion-derived measures only included the subset of controls who had DWI data available (*i.e.*, 31/57 controls). Of note, all main analyses were replicated by comparing the patients against these 31 controls (see *Supplementary Results*).

### Data preprocessing

T1w images were deobliqued, reoriented, skull stripped, and submitted to FreeSurfer (v6.0; https://surfer.nmr.mgh.havard.edu/) to extract surface models of the cortical mantle (Dale et al. 1999). Subject-specific, vertex-wise maps of cortical thickness were then generated by measuring the Euclidean distance between corresponding pial and white matter vertices.

The rs-fMRI scans were preprocessed using FSL (https://fsl.fmrib.ox.ac.uk/fsl/fslwiki/) and AFNI (https://afni.nimh.nih.gov/afni) and included: removal of the first 4 volumes from each time series to ensure magnetization equilibrium, reorientation, motion correction, skull stripping, spatial smoothing using a 6 mm FWHM Gaussian kernel, grand mean scaling, and detrending. Prior to connectivity analysis, time series were statistically corrected for effects of head motion, white matter signal, and cerebrospinal fluid (CSF) signal. They were also band-pass filtered to 0.01-0.10 Hz. Patients and controls did not differ with respect to head motion (*t*=0.84, *p*>0.20) and mean framewise displacement (*t*=1.24, *p*>0.11). Following rs-fMRI preprocessing, a boundary-based registration technique (Greve and Fischl 2009) mapped the functional time series to each participant’s cortical surface and subsequently to a 10k vertices (*i.e.*, surface points) version of the Conte69 human symmetric surface template.

Diffusion MRI data were preprocessed using MRTrix^3^ (v0.3.15; http://www.mrtrix.org/) and included correction for susceptibility distortions using FSL TOPUP (Andersson et al. 2003) as well as for motion and eddy currents using FSL EDDY. As in prior work (Larivière et al. 2019), we generated surfaces probing the white matter ~2 mm beneath the cortical interface to examine diffusion measures along the superficial white matter. Surfaces were derived by systematically contracting the white matter interface along a Laplacian potential field towards the ventricular walls. Diffusion tensor-derived fractional anisotropy (FA) and mean diffusivity (MD), surrogates of fiber architecture and tissue microstructure, were interpolated along these surfaces and mapped to Conte69.

### Distance-enriched functional connectivity analyses

Individualized functional connectomes were generated by computing pairwise correlations between all pairs of cortical surface vertices. For each region within the *z*-transformed connectome matrices, we retained the top 10% of weighted functional connections and calculated the average geodesic distance to all other regions in this connectivity profile. Distance between nodes was measured by the Chamfer algorithm which propagates geodesic distances through combined grey and white matter masks. In contrast to computing surface-based geodesic distance in each hemisphere independently (Oligschlager et al. 2017; Hong et al. 2018) the Chamfer algorithm is able to capture inter-hemispheric connections. Distances were then quantified in volume space and subsequently mapped to the surface template. The resulting connectivity distance maps recapitulate a given region’s geodesic distance to its functionally connected areas within the cortex and, as such, characterize the relationship between physical distance and functional connectivity. In contrast to conventional seed-based functional connectivity analyses, our connectivity distance profile approach provides additional topographic information that can differentiate local from distant projection patterns. Distance maps in patients were *z*-scored relative to controls and sorted into ipsilateral/contralateral to the focus. As in previous work (Bernhardt et al. 2010), surface-based linear models compared connectivity distance in patients relative to controls using SurfStat (Worsley et al. 2009), available at http://mica-mni.github.io/surfstat. Findings were corrected for age and sex, as well as for multiple comparisons at a family-wise error (FWE) rate of *p*<0.05.

*Post-hoc* analyses centered on regions of connectivity distance alterations in TLE furthermore quantified connectivity distance shifts, informed by Harrell-Davis quantile estimators (Harrell and Davis 1982). For statistical significance estimation, we computed 95% bootstrap confidence intervals of the decile-wise differences between controls and TLE, computed across 1000 iterations.

### Connectivity distance asymmetry

To assess the utility of our findings for seizure focus lateralization, we repeated the connectivity distance analysis in left and right TLE cohorts independently. Inter-hemispheric asymmetry maps of connectivity distance were also computed for each patient.

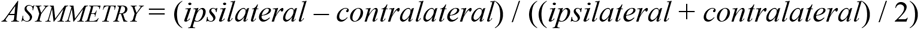

Using surface-based paired *t*-tests, we then compared ipsilateral *vs*. contralateral connectivity distance profiles in TLE.

### Rich club, community-, and gradient-based stratification

A series of analyses contextualized our findings with respect to whole-brain network topology.

#### a) Rich club

The ‘rich club’ refers to a core network consisting of high-degree and densely interconnected *hub regions* (van den Heuvel and Sporns 2011). To assess connectivity distance reductions within this specialized taxonomy, we computed the weighted rich club coefficient ɸ^w^(*k*) across the range of possible degrees and normalized it relative to 1000 randomly-generated networks with similar degree and weight properties. Nodes with degree *k* ≥ 1700 were characterized as rich club nodes and those with *k* < 1700 as peripheral. To ensure that findings were not biased by the choice of *k*, analyses were repeated across different thresholds within a ‘rich club regime’, comprising normalized ɸ^w^(*k*) values between *k* ≥ 1500 to 1900 and yielded virtually identical findings. Connections between nodes were classified into rich club (rich club– rich club), feeder (rich club–peripheral), and local (peripheral–peripheral). For each of these classes, patient-specific vertex-wise connectivity distance findings, normalized to controls, were averaged and Cohen’s *d* effect sizes were estimated.

#### b) Communities and functional gradients

We stratified distance alterations based on a functional atlas of seven resting-state networks (Yeo et al. 2011). In addition to decomposing the brain into discrete networks, the functional connectome can also be characterized by a gradient of smooth transitions running from unimodal sensory systems towards higher-order transmodal circuits, such as the default-mode and frontoparietal networks (Margulies et al. 2016). We computed the connectome gradient using 100 subjects from the Human Connectome Project and subsequently partitioned it into 20 bins, ranging from unimodal (1^st^ bin) to higher-order/transmodal regions (20^th^ bin). Mean vertex-wise connectivity distance was calculated across each community and gradient bin. Patients were then compared to controls using two-sample *t*-tests; findings were corrected for multiple comparisons using False Discovery Rate (FDR) procedures.

### Relation to cortical morphology and microstructure

To examine structural underpinnings of functional connectivity distance shifts, we repeated the above analyses while controlling for effects of cortical thickness and superficial white matter microstructure.

#### a) Cortical morphology

Surface-based linear models first compared cortical thickness in TLE compared to controls while controlling for age and sex. We then assessed functional connectivity distance alterations in TLE *vs.* controls while controlling for cortical thickness alterations at each vertex to investigate shifts in the connectivity distance distribution in TLE above and beyond effects of cortical atrophy.

#### b) Superficial white matter microstructure

The superficial white matter immediately beneath the cortex harbors termination zones of long-range tracts and short-range, cortico-cortical U-fibers (Schmahmann and Pandya 2009), and is therefore ideal to study the interplay between microstructure and functional connectivity (Larivière et al. 2019). We compared diffusion MRI-derived superficial white matter microstructure (*i.e.*, fractional anisotropy (FA) and mean diffusivity (MD), surrogates of fiber architecture and tissue microstructure) in TLE *vs.* controls. As in *a*), we compared functional connectivity distance between groups while also controlling for vertex-wise diffusion parameters to investigate functional connectivity anomalies in TLE above and beyond superficial white matter perturbations.

### Association to surgical outcome

Recent neuroimaging evidence supports an increasing role of structural and functional network modeling in informing clinical practice (Engel et al. 2013). While prior research has trained classification algorithms using isolated structural and diffusion markers to predict post-surgical seizure freedom (Bonilha et al. 2013; Gleichgerrcht et al. 2018), the utility of functional connectivity in this context remains underexplored. Here, we leveraged a supervised pattern learning paradigm to evaluate whether functional connectivity distance shifts can predict post-surgical seizure outcome in individual patients. Specifically, we build learners discriminating seizure-free (Engel I) from non-seizure-free (Engel II-IV) patients. To minimize overfitting, a two-step dimensionality reduction was performed on individual-level connectivity distance maps. Maps were first parcellated using the Desikan-Killiany atlas, yielding 68 features for each individual map, and subsequently fed into a principal component analysis to generate orthogonal components. These components were submitted to a L2-regularized logistic regression models. Training and performance evaluation employed nested 5-fold cross-validation with 100 iterations to allow for unbiased and conservative performance assessment of previously unseen cases. Statistical performance of the model was assessed using 1000 permutation tests with randomly shuffled surgical outcome labels. The combination of PCA dimensionality reduction and L2 regularization, together with the use conservative cross-validation techniques were chosen to reduce overfitting of the classification. All methods were implemented using the openly-available Scikit-learn package for Python (Abraham et al. 2014). To index feature importance across cortical regions, we projected the product of logistic regression coefficients and principal component scores onto the cortical surface, with positive feature weight values predicting seizure-free outcome and negative values predicting non-seizure-free outcome.

To combine pre-operative connectivity distance feature data with surgical information, we related the classifier weights to surgical resection cavities in individual patients. We automatically segmented post-surgical cavities by registering both pre- and post-operative T1w images to the MNI152 standard template through linear transformations and subsequently subtracting the post-operative scan from the pre-operative scan. Segmented cavities were visually inspected and manually edited to ensure that the extent of the resections was correctly identified. Patient-specific cavities were then mapped and registered to the surface template, and a consensus label was generated as defined by the union of all segmentations. To associate resection information with the statistical pattern learning results, we computed classifier weights within and outside the patient-specific cavity masks.

To assess whether functional connectivity distance features can improve prediction of post-surgical seizure outcome compared to standard clinical and neuroimaging parameters, we benchmarked classification performance of our surface-based predictive model against a model based on socio-demographic and clinical variables (age, sex, duration of epilepsy, hippocampal volume ipsilateral and contralateral to the seizure focus). Both classifiers were compared using McNemar’s tests, as implemented in the Python package Statsmodels.

## Results

### Connectivity distance reductions in TLE

In controls, we observed inter-regional variations in average connectivity distance across the cortex, with transmodal networks showing longest distances and sensory cortices showing shortest distances (Figure 1A). Notably, TLE patients demonstrated marked reductions relative to controls (Figure 1B) in a bilateral network encompassing lateral temporo-limbic cortices (ipsilateral: *p*_*FWE*_<0.001; contralateral: *p*_*FWE*_=0.05) and dorsomedial prefrontal regions (ipsilateral: *p*_*FWE*_=0.09; contralateral: *p*_*FWE*_<0.005), as well as contralaterally in ventromedial prefrontal cortex (*p*_*FWE*_<0.05) and ipsilaterally in inferior frontal gyrus (*p*_*FWE*_=0.06).

**Figure 1.**
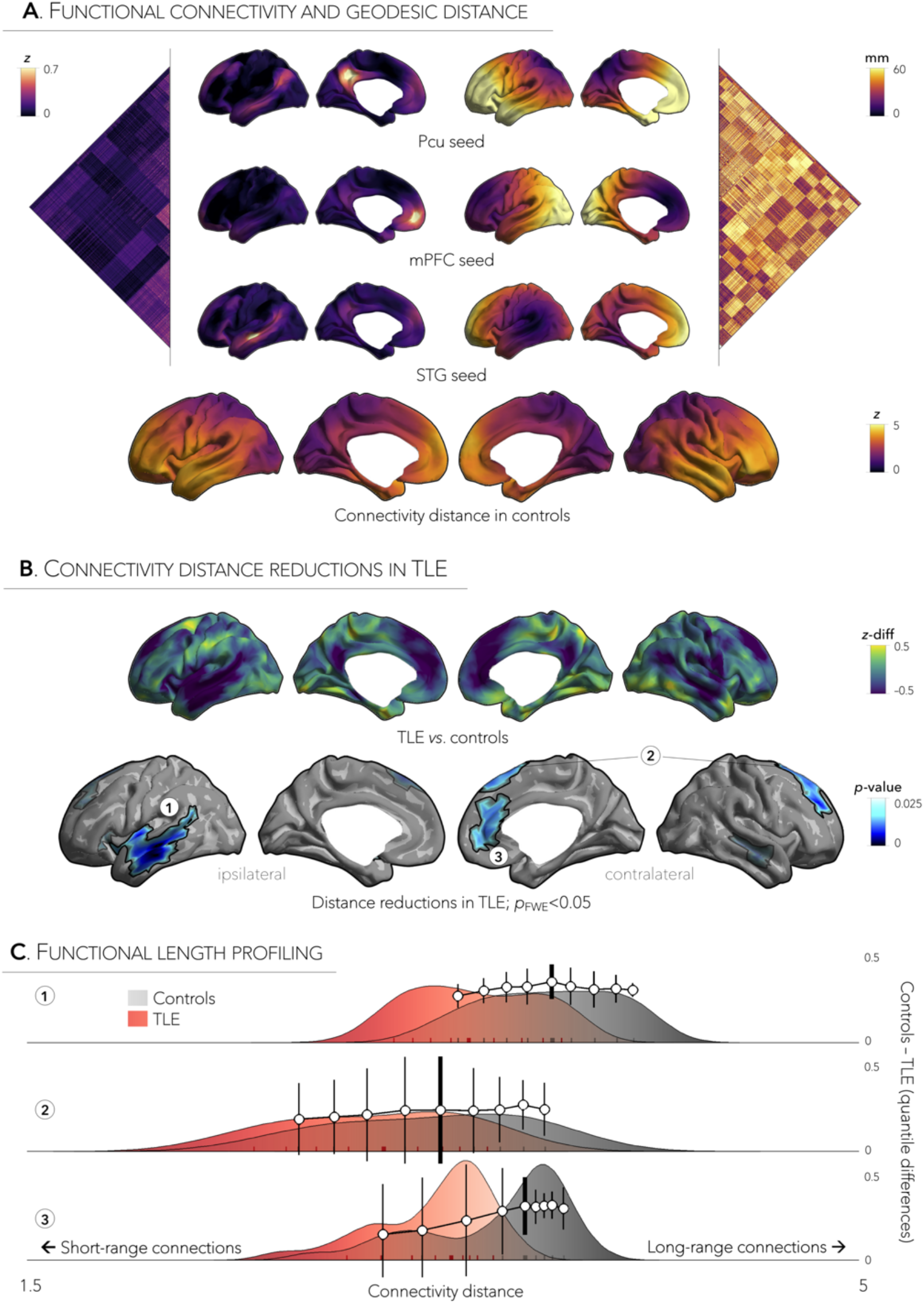
Connectivity distance reductions in temporal lobe epilepsy (TLE). a | Functional connectivity and geodesic distance profiles are displayed for three exemplary regions: precuneus (Pcu; *top*), medial prefrontal cortex (mPFC; *middle*), and superior temporal gyrus (STG; *bottom*). Cortex-wide variations in connectivity distance in healthy controls, with transmodal networks showing longest distances and sensory and motor cortices the shortest. b | Relative to controls, TLE patients showed marked connectivity distance reductions in bilateral temporal and insular cortices and dorsomedial prefrontal regions, as well as contralateral ventromedial prefrontal cortex, and ipsilateral inferior frontal gyrus. Significant clusters are numbered 1–3. Trends are shown in semi-transparent. c | Connectivity distance reductions in these clusters were driven by concurrent increases in short-range and decreases in long-range connections in TLE. Shift functions are superimposed onto each distribution and illustrate by how much connectivity distance profiles are shifted in TLE (red) relative to controls (grey). Vertical lines along this function indicate the 95% bootstrap confidence interval for each decile difference (Controls – TLE). Shorter lines at the bottom of each distribution mark the deciles for each group; thicker lines represent the median.

Density distribution of every vertex within these three significant clusters, sorted by the average geodesic distance of that vertex to its functionally connected areas, indicated that connectivity distance changes related to concurrent increases in short-range and decreases in long-range connections in TLE (Figure 1C).

### Relation to seizure focus laterality

Dominant patterns of connectivity distance reductions involving the ipsilateral temporal cortex were observed in left and right TLE cohorts independently (*p*_*FWE*_<0.001, Figure 2A). Similarly, analysis of within-patient asymmetry maps revealed pronounced connectivity distance reductions in ipsilateral temporal regions (*p*_*FWE*_<0.001, Figure 2B). Individual analyses in this cluster confirmed that the majority of patients (23/30; 77%) showed dominant ipsilateral, compared to contralateral, reductions.

**Figure 2.**
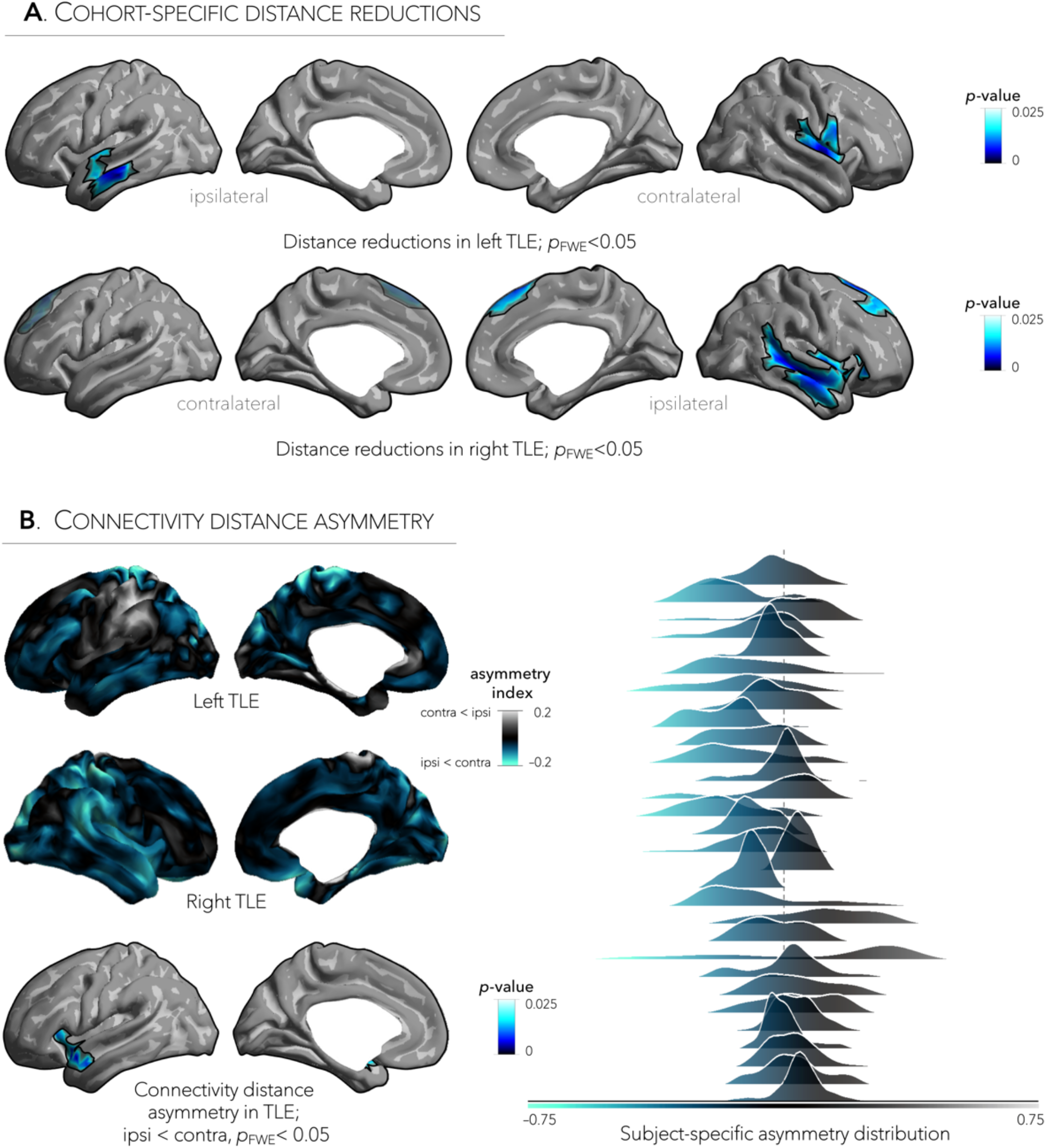
Connectivity distance asymmetry as a lateralizing sign in TLE. **a |** Dominant patterns of connectivity distance reductions in left and right TLE encompassed an ipsilateral network involving temporal and insular cortices. Trends are shown in semi-transparent. **b |** Asymmetry maps comparing ipsilateral *vs*. contralateral hemispheres in left and right TLE cohorts showed connectivity distance reductions in ipsilateral anterior temporal regions, and subject-specific asymmetry values within this cluster showed consistent asymmetry in 77% of patients.

### Relation to connectome topology

Considering key features of large-scale network organization, strongest connectivity distance reductions in TLE were observed among rich club nodes (Figure 3A), indicating higher susceptibility of hub regions to connectivity contractions. Similarly, community-based stratification (Yeo et al. 2011) and connectome topographic profiling (Margulies et al. 2016) showed strongest effects in the default-mode network (*p*_*FDR*_<0.025, Figure 3B), a network situated at the transmodal apex of the adult connectome gradient (*p*_*FDR*_<0.025, Figure 3C).

**Figure 3.**
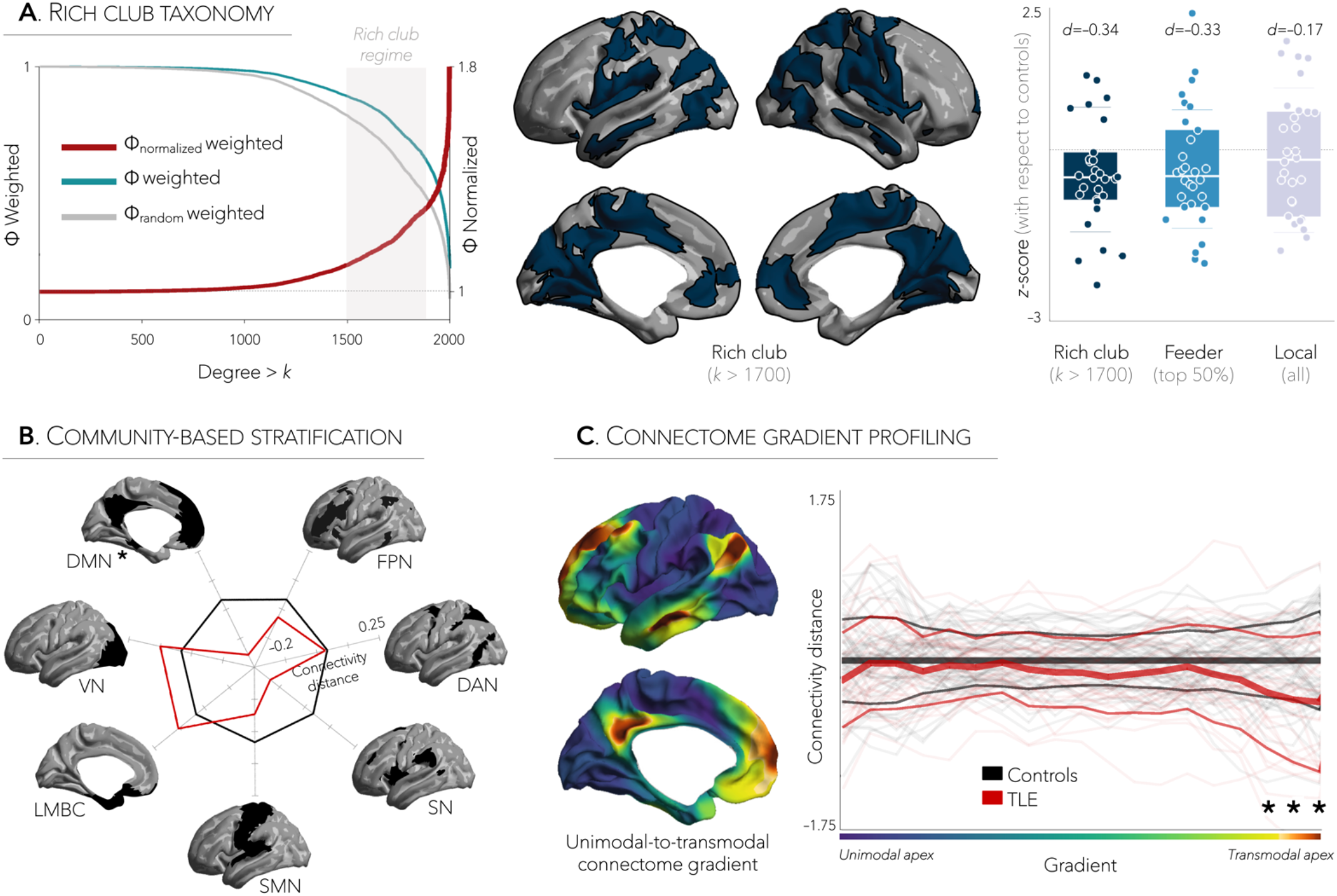
Connectivity distance reductions in TLE co-localize with high-degree, transmodal networks. **a |** The distribution of rich club nodes was mapped onto the cortical surface and Cohen’s *d* effect sizes indicated that connectivity distance reductions co-localized with rich club (rich club–rich club) and feeder (rich club–local) connections. **b |** Stratification of the functional connectome into seven canonical resting-state networks revealed that TLE patients have preferential connectivity distance reductions in default-mode (DMN) network. **c |** Similarly, reductions in TLE closely co-localized with the transmodal apex of the adult connectome gradient, which differentiates between unimodal, sensory and higher-order, transmodal systems. * = *p*_*FDR*_<0.025.

### Connectivity reductions are mediated by microstructural damage

Compared to controls, patients presented with cortical thinning across bilateral premotor, superior frontal, and paracentral cortices (*p*_*FWE*_<0.001), ipsilateral temporal pole extending into mesiotemporal regions (*p*_*FWE*_<0.001, Figure 4B). Spatial comparison between areas of cortical atrophy and patterns of connectivity distance reductions revealed minimal overlap (*Dice*=0.02); and findings of connectivity reductions remained robust after correcting for atrophy, suggesting independence from morphological alterations.

**Figure 4.**
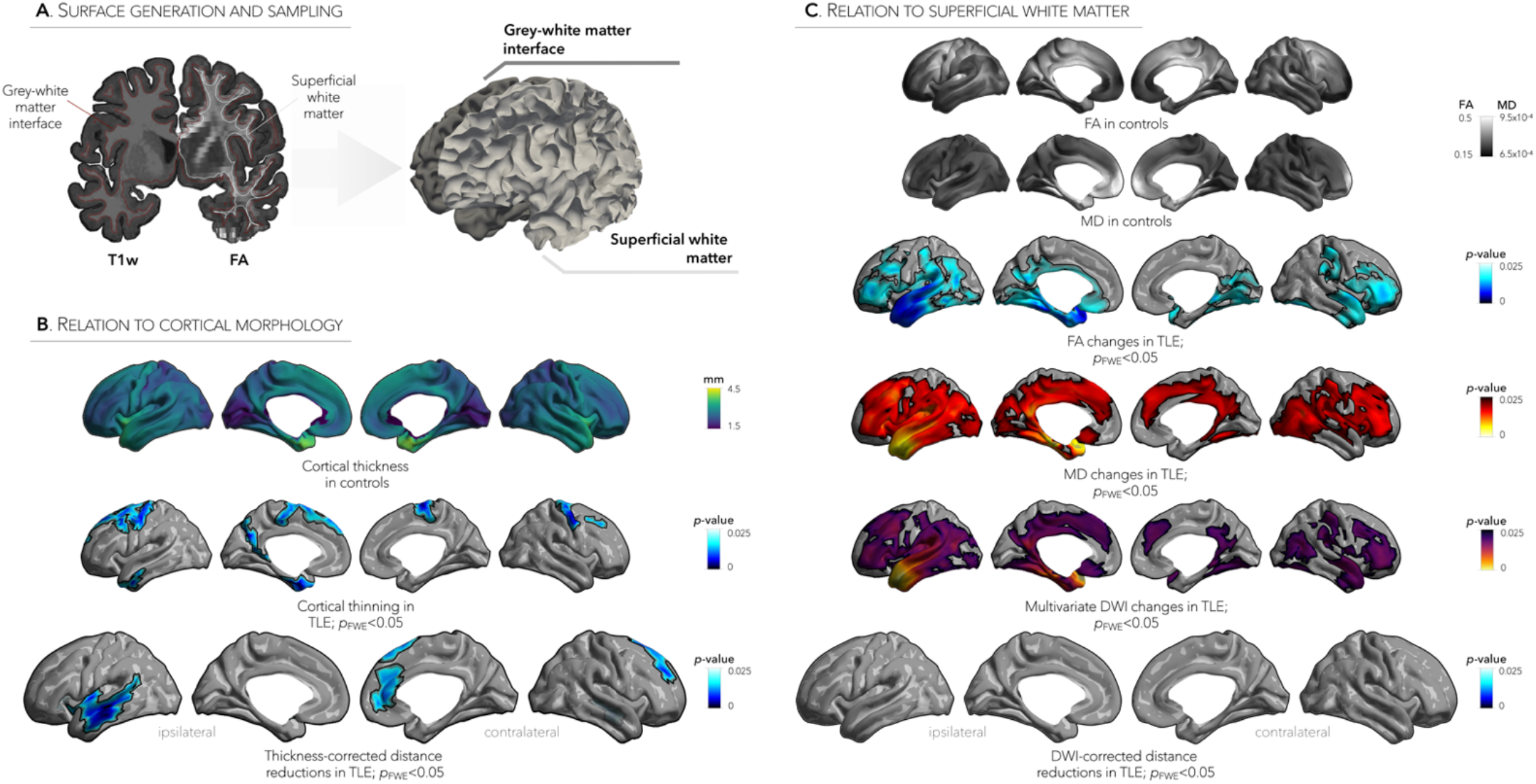
Microstructural damage mediates connectivity distance reductions. **a |** Cortical thickness was measured at each vertex as the Euclidean distance between white and pial surfaces, whereas fractional anisotropy (FA) and mean diffusivity (MD) were sampled along superficial white matter surfaces generated 2 mm below the grey-white matter interface. **b |** Patterns of cortical thinning differed from that of functional connectivity distance reductions in TLE, suggesting that shifts in connectivity distance occurred independently of cortical atrophy. **c |** Repeating the connectivity distance analysis while controlling for superficial white matter effects revealed no significant changes in TLE compared to controls, suggesting a mediatory role of microstructural damage in functional distance reductions.

We also observed extensive superficial white matter diffusion alterations in TLE relative to controls, albeit in a different spatial distribution as cortical atrophy. Microstructural alterations, characterized by reduced FA and increased MD, were located in the ipsilateral temporal pole (*p*_*FWE*_<0.001), and extended centrifugally to bilateral frontal, temporal, and occipital cortices (*p*_*FWE*_<0.05, Figure 4C). Overlaps with functional changes were modest (*Dice*=0.12). Moreover, the effect size of connectivity distance reductions in TLE was markedly reduced when repeating analyses while controlling for superficial white matter metrics (effect size reductions up to 45% across clusters of findings; with strongest reductions in ipsilateral temporal regions), suggesting a mediatory role of microstructural damage in functional contractions.

### Associations to post-surgical seizure recurrence

A supervised pattern learning paradigm evaluated whether post-surgical seizure freedom imparts distinct signatures on connectivity distance profiles. Dimensionality reduction of the connectivity distance maps yielded 24 principal components explaining 98.7% in variance. Using 5-fold cross-validations, we found that connectivity distance predicted surgical outcome (seizure-free *vs*. non-seizure-free) at a mean±SD accuracy of 74±8% (100 iterations). Permutation tests with 1000 randomly shuffled outcome labels indicated that performance exceeded chance levels (*p*<0.05, Figure 5A).

**Figure 5.**
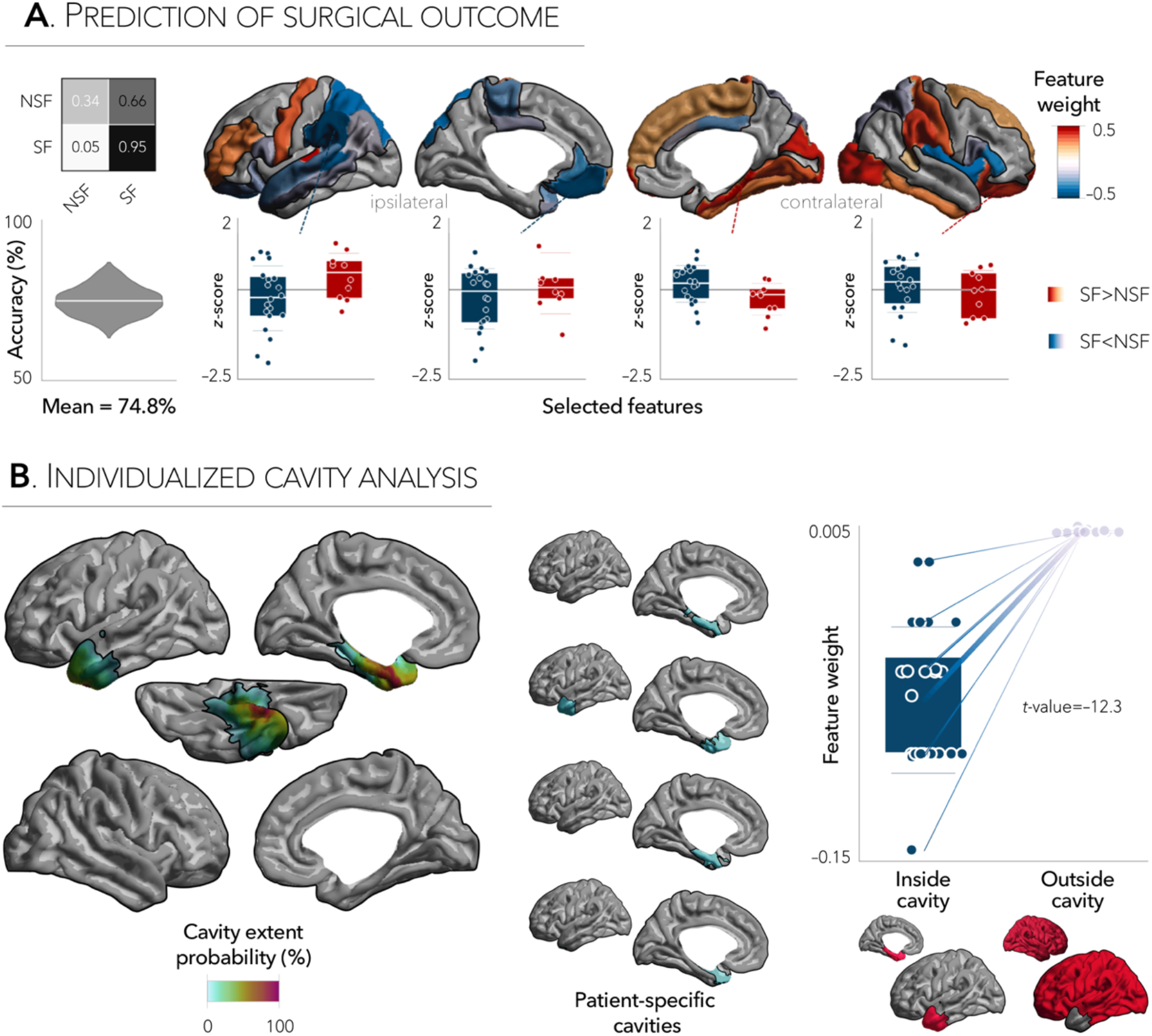
Connectivity distance signature of surgical outcome. **a |** A supervised pattern learning algorithm was trained on functional connectivity distance data to predict surgical outcome in patients (seizure-free; SF *vs*. non-seizure-free; NSF) and achieved 74.8% accuracy. Confusion matrix summarizing the model’s performance is displayed in inset (values range from 0 to 1). Most important features (top and bottom 25%) are mapped to the cortical surface while boxplots for four exemplary regions depict mean connectivity distance values as a function of group. **b** | Cavity extent probability across patients, and individual cavity maps for a subset of patients are projected onto the surface template. Seizure free patients showed more marked connectivity distance reductions in ipsilateral regions, whereas those with persistent seizures were characterized by connectivity distance reductions in regions contralateral to the surgical cavity.

Features associated with seizure freedom included connectivity distance reductions in ipsilateral default-mode regions and contralateral temporo-insular cortices. Conversely, functional contractions associated with seizure recurrence were predominantly located contralaterally in mesial temporal, orbitofrontal, visual, and centro-parietal cortices, as well as in ipsilateral frontopolar cortex (Figure 5A). Stratification of classifier weights inside/outside the patient-specific resection cavities indicated that connectivity distance reductions inside the cavity were associated with seizure freedom, while reductions outside the cavity, specifically in contralateral regions, predicted seizure recurrence. This shift was consistent across patients and highly significant (paired *t*-test, p<0.001; Figure 5B).

Our new predictive model based on functional connectivity distance features performed significantly better than a model based on socio-demographic and clinical variables (age, sex, hippocampal volumes, duration of epilepsy). Indeed, compared to our model, the latter had a lower mean±SD accuracy of 61.3±5% (using 5-fold cross-validation and 100 iterations). Direct comparisons of both classifiers using McNemar’s tests revealed that prediction models based on connectivity distance features outperformed those based on clinical and hippocampal variables (chi-square=44.4, *p*<0.0001).

## Discussion

Our macroscale connectivity analysis allowed us to interrogate mechanisms suggested by experimental models of limbic epilepsy, where connectivity contractions may relate to the formation of topologically isolated excitatory circuits that contribute to recurrent seizures (Knopp et al. 2005; Sharma et al. 2007), in a cohort of surgical candidates with TLE. Notably, our analysis was centered on a novel *in vivo* approach that combined resting-state functional connectivity and physical distance between cortical areas, and which allowed us to parameterize the balance between short- and long-range functional connections in TLE. Comparing drug-resistant TLE patients to controls, we observed marked reductions in temporo-limbic and dorsomedial prefrontal cortices, connectivity contractions which were found to be driven by concomitant increases in short-range and decreases in long-range connections in TLE. Exploring co-occurring morphological and microstructural perturbations, we observed that connectivity reductions were independent of temporal and fronto-central cortical thinning but mediated by superficial white matter alterations stemming from the mesiotemporal epicenter. A supervised machine learning algorithm with conservative 5-fold cross-validation further identified salient connectivity distance features that predicted post-surgical seizure recurrence with 75% accuracy and outperformed learners based on clinical and hippocampal MRI features, suggesting prognostic utility. Collectively, our findings provide new *in vivo* evidence for topological isolation as a potential disease mechanism of TLE, and support promising clinical benefits of combining connectomic data with physically-grounded information.

Our study harnessed rs-fMRI, a non-invasive window to probe intrinsic functional networks in a highly reproducible and individualized manner. As opposed to conventional rs-fMRI connectivity analyses, which focus on *a-priori* defined seeds or discrete communities, our novel framework enables the profiling of connectivity distances along the cortical sheet. This surface-based approach benefits from a continuous, and high-resolution reference frame to assess the interplay between functional architecture, cortical morphology, and white matter microstructure. Mirroring previously reported maps of cortical distance in healthy individuals (Oligschlager et al. 2017), we observed marked inter-areal variations in the average physical distance of a region’s functional connections. Interestingly, when compared to controls, TLE patients presented with a significant loss of long-range communication pathways emanating from bilateral temporo-insular cortices, a finding consistent with covariance analyses focused on the mesiotemporal circuitry showing topologically isolated networks in drug-resistant TLE (Bernhardt, Bernasconi, Hong, et al. 2016). Importantly, assessing left and right TLE patients separately yielded dominant ipsilateral temporo-limbic effects and, as such, confirmed robustness of our findings. At the individual level, connectivity distance reductions restricted to the anterior temporal cortex showed consistent asymmetry in 77% of patients, supporting potential clinical benefits of our approach for focus lateralization. The co-localization of functional connectivity anomalies with the mesiotemporal epicenter motivates future work to examine associations between our imaging-derived connectome measures and neurophysiological markers that may index epileptogenicity, notably abnormal rates of high-frequency oscillations and content of low-voltage fast activity (Bartolomei et al. 2008; Frauscher et al. 2018).

By mapping the connectivity distance of every region along the cortical landscape, our framework reconciled functional organizational features with MRI substrates of cortical morphology and microstructure. Cortical atrophy in TLE patients followed a bilateral temporal and fronto-central spatial topography, a finding in agreement with previous cross-sectional (Lin et al. 2007; McDonald et al. 2008; Bernhardt et al. 2010) and longitudinal imaging studies (Bernhardt et al. 2009; Galovic et al. 2019), along with a recent multi-site meta-analysis (Whelan et al. 2018). In contrast to widespread cortical atrophy, superficial white matter perturbations (Liu et al. 2016) followed a specific temporo-limbic spatial pattern with dominant effects ipsilateral to the focus. Previous studies relating structure and function in the healthy connectome have generally yielded stronger correspondence of functional interactions with structural connectivity estimated from diffusion imaging than with T1w-derived estimates of cortical morphology (Misic et al. 2015). Several lines of evidence support the interpretation that widespread cortical thinning in TLE may be more closely related to consequences of the disease or from living with epilepsy. These findings are supported by previous cross-sectional and longitudinal studies showing greater atrophy in hippocampal, neocortical, and subcortical regions in patients with a longer disease duration or those with a higher seizure burden (Bernhardt et al. 2009; Coan et al. 2009; Caciagli et al. 2017; Galovic et al. 2019), as well as studies showing effects of medication load on measures of cortical morphology (Pardoe et al. 2013). On the other hand, our functional connectivity findings strongly resembled superficial white matter changes in the temporo-limbic circuitry, supporting a potential mediatory role of this compartment on functional anomalies in drug-resistant TLE. In addition to their spatial resemblance to the agranular/dysgranuar limbic territory, white matter changes have also been more consistently related to the laterality and severity of hippocampal pathology than grey matter cortical thinning, and may therefore reflect cascading effects secondary to mesiotemporal pathology in TLE (Tavakol et al. 2019). Such findings were also echoed in diffusion MRI studies that profiled deep white fiber tracts (Concha et al. 2012; Keller et al. 2015), showing anomalies tapering off with increasing distance from the affected mesiotemporal lobe. By consolidating previously reported disease effects from multiple spatial scales with functional network profiles, our study offers non-invasive insights into the mechanisms by which mesiotemporal anomalies may ultimately propagate to whole-brain functional networks in TLE (Lariviere et al. 2019).

Our surface-based findings, particularly the reductions in long-range connectivity together with increases in local connections in temporo-limbic circuits may provide a physical mechanism underlying previous graph theoretical findings obtained from various connectivity modalities (van Diessen et al. 2014). These changes were previously labelled as network regularization, and frequently observed in experimental models of cortical epilepsy (Otte et al. 2012) as well as in patients with drug-resistant focal epilepsy (van Diessen et al. 2014). While several network configurations may promote spontaneous seizures (Jirsa et al. 2014), spatial compactness of local circuits has been suggested as a plausible mechanism that tilts the balance towards epileptogenesis by facilitating recurrent activity and local oscillations (Sharma et al. 2007). Prior EEG/SEEG studies have indeed reported network regularization at seizure onset, a configuration that shifts towards a globally integrated process as the seizure progresses, eventually reaching a random configuration upon seizure termination (Schindler et al. 2008). To further explore computational mechanisms of seizure dynamics, it has been speculated that local clusters of neurons may isolate themselves from the rest of the network, likely through an imbalance between local and distant connections, a mechanism described as "network tightening" (Khambhati et al. 2015). Under this account, functional connectome contractions in TLE may be speculated to play a role in annihilating epileptogenic activity and, as such, may represent a potential compensatory mechanism through which locally confined, seizure-related activity is prevented from spreading.

We close by highlighting that our measures were ~75% accurate in predicting post-surgical seizure outcome, outperforming learners based on clinical and hippocampal volumetric features. Dominant connectivity distance reductions in ipsilateral regions related to better outcomes, while signatures predicting seizure recurrence encompassed extra-temporal and contralateral regions, a finding that parallels predictors of surgical failure seen at the level of morphological (Bernhardt et al. 2015) and diffusion MRI measures (Gleichgerrcht et al. 2018). Findings could be further contextualized in light of our inclusion criteria, which required post-surgical MRI and histological confirmation of the resected cavity in all patients. Indeed, overlapping patient-specific cavity data with machine learning signatures revealed that the balance of anomalies inside *vs*. outside the cavity served as a key driver for outcome prognosis. Additional strengths of our approach rely on the use of conservative cross-validation for training and validation as well as relatively long post-surgical follow-up times averaging three years. Nevertheless, in light of prior work suggesting the possibility of seizure recurrence at longer follow-up times in post-surgical patients that were initially considered seizure-free (de Tisi et al. 2011), we cannot rule out potential sources of uncertainty in the surgical outcome labels used in our study. Future studies may also better characterize the generalizability and predictive power of our findings in other prevalent drug-resistant focal epilepsies related to malformations of cortical development, notably focal cortical dysplasia with subtle structural imaging signs. Lastly, large-scale initiatives such as ENIGMA-epilepsy (Whelan et al. 2018) have demonstrated that it is possible to aggregate and coordinate neuroimaging data, clinical information, and analytical strategies across sites. In this context, our surface-based features are openly available to the scientific community, and we hope that similar efforts will help with the validation and dissemination of novel biomarkers.

## Acknowledgements

SL acknowledges funding from Fonds de la Recherche du Québec – Santé (FRQ-S) and the Canadian Institutes of Health Research (CIHR). RvdW receives support from a Savoy Foundation studentship. BF receives funding from FRQ-S (Chercheur-Boursier clinician Junior 2). AB and NB were supported by FRQ-S and CIHR (MOP-57840, MOP-123520). ZZ was supported by National Science Foundation of China (NSFC: 81422022; 863 project: 2014BAI04B05 and 2015AA020505) and China Postdoctoral Science Foundation (2016M603064). BCB acknowledges research funding from the SickKids Foundation (NI17-039), the National Sciences and Engineering Research Council of Canada (NSERC; Discovery-1304413), CIHR (FDN-154298), Azrieli Center for Autism Research (ACAR), an MNI-Cambridge collaboration grant, and the Canada Research Chairs (CRC) Program.

