## Supplementary Materials for "Functional connectome contractions in temporal lobe epilepsy: microstructural underpinnings and associations to surgical outcome"

[1]*Multimodal Imaging and Connectome Analysis Laboratory, McConnell Brain Imaging Centre, Montreal Neurological Institute and Hospital, McGill University, Montreal, QC, Canada;* [2]*Department of Medical Imaging, Jinling Hospital, Nanjing University School of Medicine, Nanjing, China;* [3]*Montreal Neurological Institute and Hospital, McGill University, Montreal, QC, Canada;* [4]*Department of Radiology, Nanjing Drum Tower Hospital, The Affiliated Hospital of Nanjing University Medical School, Nanjing, China;* [5]*NeuroImaging of Epilepsy Laboratory, McConnell Brain Imaging Centre, Montreal Neurological Institute and Hospital, McGill University, Montreal, QC, Canada;* [6]*BC Children's Hospital, Department of Pediatrics, University of British Columbia, Vancouver, BC, Canada*

[*] *authors contributed equally*

### SUPPLEMENTARY MATERIALS

**CORRESPONDENCE TO:**
Boris C. Bernhardt, PhD
Montreal Neurological Institute (NW-256)
3801 University Street
Montreal, Quebec, Canada H3A 2B4


Sara Larivière, PhD Candidate


### SUPPLEMENTARY RESULTS

#### *Sensitivity analyses*

As only a subset of control participants had DWI data available (*i.e.*, 31/57), we performed several sensitivity analyses to ensure reproducibility of our functional connectivity distance findings. Accordingly, we repeated the above analyses comparing the patients to the 31 of controls that underwent diffusion MRI, and observed virtually identical findings as in our main analyses. **(FIGURES S1, S2)**.

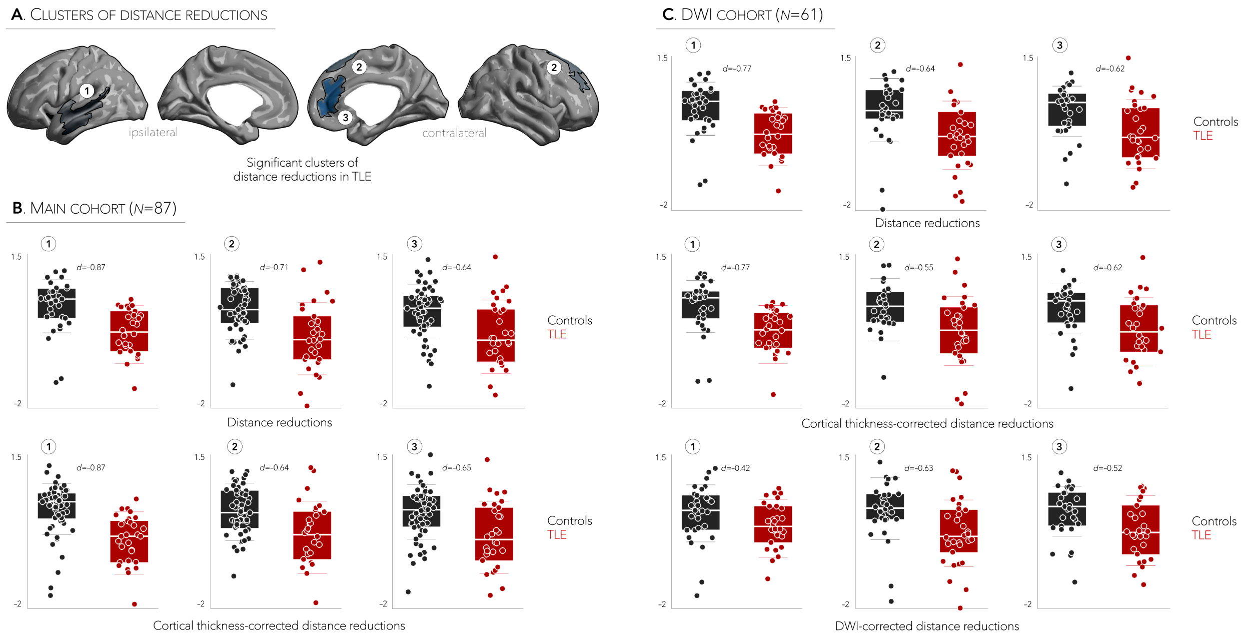


**FIGURE S1**. **Effects were robust across main and DWI-only cohorts**. **a |** Clusters of significant connectivity distance reductions in TLE patients, relative to controls, are mapped to the surface template. Cohen's *d* values measuring effect sizes for each cluster in **b |** the main cohort and **c |** a subset of participants who underwent additional diffusion MRI.

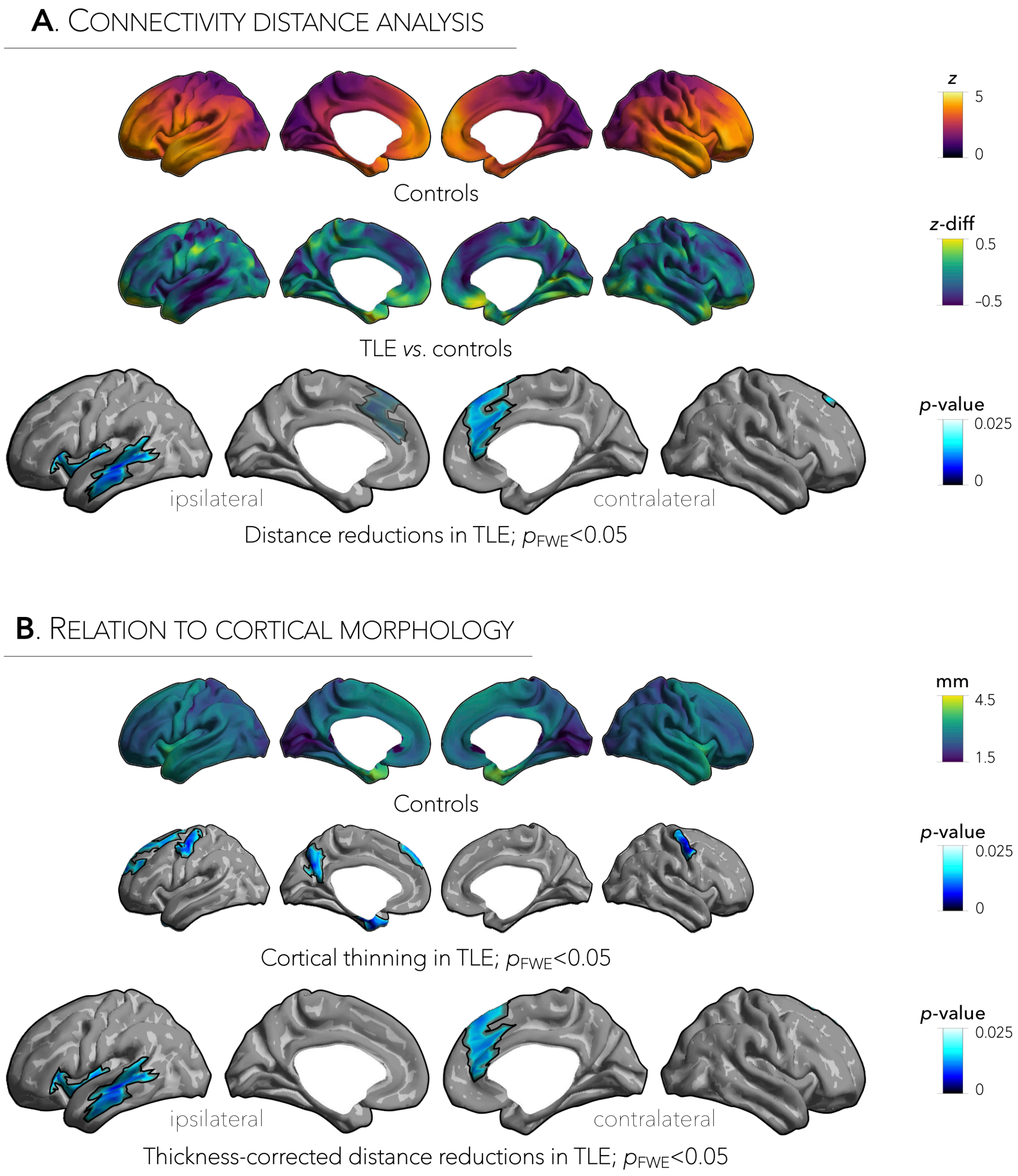


**FIGURE S2. Connectivity distance reductions in TLE are replicable in the DWI-only cohort.** **a |** Connectivity distance reductions were reproduced in a subset of participants who underwent additional diffusion MRI ($n_{\text{controls}}$=31/57, $n_{\text{TLE}}$=30/30) with strongest reduction effects observed in ipsilateral temporo-insular cortices ($p_{FWE}$<0.01) and bilateral superior frontal regions (ipsilateral: $p_{FWE}$=0.06; contralateral: $p_{FWE}$<0.05). **b |** Cortical atrophy in patients encompassed bilateral fronto-central and temporal regions ($p_{FWE}$<0.005), and was independent of connectivity distance reductions ($p_{FWE}$<0.01).

**SUPPLEMENTARY TABLE 1**. Patient-specific clinical information.

| **Patient** (side) | **Age** (years) | **Sex** | **Epilepsy duration** (months) | **Post-op. follow-up** (years) | **Surgical outcome** |
|---|---|---|---|---|---|
| 1 (R) | 25 | F | 7 | 5 | 3 |
| 2 (R) | 17 | M | 146 | 7 | 1 |
| 3 (R) | 18 | M | 213 | 2 | 2 |
| 4 (R) | 26 | F | 194 | 1 | 1 |
| 5 (R) | 26 | M | 31 | 2 | 1 |
| 6 (R) | 37 | F | 201 | 1 | 3 |
| 7 (R) | 17 | M | 67 | 7 | 1 |
| 8 (R) | 29 | F | 252 | 1 | 1 |
| 9 (R) | 36 | M | 141 | 3 | 1 |
| 10 (R) | 37 | M | 248 | 7 | 1 |
| 11 (R) | 22 | F | 37 | 6 | 1 |
| 12 (R) | 31 | F | 74 | 3 | 1 |
| 13 (R) | 37 | M | 333 | 3 | 1 |
| 14 (R) | 15 | M | 104 | 3 | 2 |
| 15 (R) | 25 | F | 53 | 3 | 1 |
| 16 (R) | 27 | F | 72 | 3 | 1 |
| 17 (R) | 16 | F | 145 | 2 | 4 |
| 18 (R) | 43 | M | 215 | 1 | 1 |
| 19 (R) | 36 | M | 2 | 2 | 1 |
| 20 (L) | 25 | F | 144 | 1 | 1 |
| 21 (L) | 23 | F | 199 | 2 | 3 |
| 22 (L) | 18 | M | 190 | 1 | 1 |
| 23 (L) | 17 | F | 225 | 2 | 3 |
| 24 (L) | 19 | F | 128 | 5 | 1 |
| 25 (L) | 23 | M | 84 | 9 | 1 |
| 26 (L) | 37 | M | 356 | 1 | 1 |
| 27 (L) | 25 | M | 132 | 8 | 2 |
| 28 (L) | 30 | F | 77 | 1 | 1 |
| 29 (L) | 22 | M | 3 | 2 | 2 |
| 30 (L) | 40 | F | 442 | 3 | 4 |
